# No group-level changes in corticospinal excitability following low-intensity theta-burst ultrasound stimulation of the primary motor hand area

**DOI:** 10.64898/2026.01.06.697894

**Authors:** J Laugesen, S Bertino, MM Beck, B Sigurðsson, A Thielscher, L Christiansen, HR Siebner

## Abstract

**Background:** Low-intensity transcranial ultrasound stimulation (TUS) is a novel non-invasive brain-stimulation technique offering high spatial precision and depth penetrance. Theta burst TUS (tbTUS), featuring a temporal theta burst stimulation pattern, has been reported to facilitate corticomotor excitability, when applied to the human primary motor hand area (M1-HAND). However, replication attempts have yielded inconsistent results.

**Methods:** Fifteen healthy human participants underwent neuronavigated tbTUS targeting the left M1-HAND in the anterior wall of the precentral sulcus, as well as anterior and posterior active control sites. The three tbTUS conditions were applied in counterbalanced order on separate days. Corticospinal excitability was assessed via motor evoked potentials (MEPs) recorded before and 5, 15, 30, and 60 minutes after tbTUS, and analyzed using an rmANOVA.

**Results:** None of the tbTUS conditions produced consistent group-level changes in MEP amplitude at any time point. Both intra- and inter-subject variability were high, and individual MEP changes following precentral tbTUS did not correlate with changes after control-site stimulation.

**Conclusions:** We did not observe reliable modulatory effects of neuronavigated tbTUS on corticospinal excitability. Methodological and hardware differences may account for discrepancies across studies. Our findings align with recent reports questioning the robustness of tbTUS-induced facilitation.

## Introduction

Non-invasive transcranial brain stimulation (NTBS) techniques like transcranial magnetic and electric stimulation (TMS and TES) are established powerful tools for both neuromodulation and treatment (Wassermann et al., 2024). Yet, both techniques suffer from limited focality and limited depth penetration (Siebner et al., 2022. Paulus, 2014). Low intensity transcranial focused ultrasound stimulation (TUS) is a relatively new technique that offers greater focality and depth penetration (Murphy et al., 2024). Through the delivery of focused acoustic pressure waves, TUS is thought to influence neuronal excitability via mechanical interactions with ion channels embedded in cell membranes. Making TUS a promising approach to selectively target deep brain areas (Darmani et al., 2025), however selective targeting of well-defined cortical patches is also possible.

Repetitive TMS (rTMS) is widely used to induce neuroplasticity in research and therapeutic settings (Regengold et al., 2022., Ziemann et al., 2008., Suppa et al., 2022.). The mechanisms of rTMS-induced neuroplasticity have been mainly explored targeting the primary motor hand area (M1-HAND) (Hallett, 2007), because this region can be easily stimulated with TMS and aftereffects can be readily measured by recording the motor-evoked potentials (MEPs) in the contralateral hand. Several rTMS protocols have been introduced that can induce long-lasting effects in corticospinal excitability. with the direction of the induced plasticity (e.g. potentiation, depotentiation) depending on the temporal pattern of TMS pulses, albeit there is substantial inter- and intra-individual variability (Ziemann et al., 2008. Ziemann & Siebner, 2015., Hamada et al., 2013.). 5Hz rTMS trains and continuous theta burst stimulation, consisting of 50 Hz three-pulse bursts repeated at an inter-burst rate of 5Hz) have been shown to increase and decrease MEP amplitude, respectively (Ziemann et al., 2008).

These rTMS protocols have inspired researchers to mimic the temporal structure with TUS. Mimicking the continuous theta burst stimulation (cTBS) protocol that had been introduced by Huang et al. (2005), Zeng et al (2022) recently introduced a TUS protocol consisting of a 20ms burst of ultrasound repeated at 5Hz referred to as theta burst TUS (tbTUS).

While the original TMS-based cTBS protocol reduced corticospinal excitability, the tbTUS protocol consistently increased corticospinal excitability when applied to the M1-HAND (Zeng et al., 2022). The facilitatory effect of tbTUS has been replicated multiple times by the same group (Shamli Oghli et al., 2023. Samuel et al., 2023., Ding et al., 2024), while others found opposite effects of tbTUS (Bao et al., 2024) or or no consistent modulation of corticospoinal excitability (Fong et al., 2024).

In the present study, we reinvestigated the after-effects of tbTUS targeting the M1-HAND on corticospinal excitability. We specifically targeted the anterior wall of the central sulcus, as it hosts the large corticomotoneuronal cells thought to convey the TMS-evoked activity to the spinal motoneurons (Siebner et al., 2022). to test spatial specificity, tbTUS was delivered to two active control sites placed anteriorly and posteriorly to the verum target in two separate experimental sessions.

## Methods and materials

### Participants

Fifteen healthy volunteer participants (nine females, age range: 23-28 years) were recruited for the study. We estimated our required sample size using values reported in Zeng et al (2022) (43.1% ± 25.1% SD change in MEP amplitude from before vs after tBTUS compared to sham). This yielded an effect size of approx. dz = 1.7. This effect would be 99% powered at N=10 but expecting more subtle effects due to using active controls, we opted to test 15 subjects. Participants had no history of neuropsychiatric diseases or use of psychoactive medications. Contraindications to TMS and MR were excluded at inclusion. Possible exclusion criteria were history of epilepsy or familiar history of epilepsy, neurological or neuropsychiatric disorders, for brain stimulation, and claustrophobia, pregnancy or implanted metals, for MR, among others. No participants reported any side effects. The study was conducted in accordance with the 8th Declaration of Helsinki and all participants gave written informed consent before the experiments.

### Experimental procedure

#### Magnetic resonance imaging

Before experimental sessions, a whole-brain volumetric structural MRI scan was acquired using a T1-weighted sequence (voxel size 0.9x0.9x0.9 mm, slice thickness 0.9 mm, repetition time (TR) 2700 ms, echo time (TE) 3.7 ms, inversion time (TI) 1090 ms, flip angle 9°, field-of-view (FOV) 260x260mm). MRI images were acquired on a Siemens Prisma 3T scanner using a 32-channel receive head coil (Siemens, Erlangen, Germany). Four participants had pre-existing T1-weighted brain scans that had been acquired as part of other studies, acquired using the same scanner and the same acquisition parameters and consented to reuse of these scans.

#### Neuronavigation

To secure consistent placement, positioning of the TMS coil and TUS transducer were navigated using the individual brain MRI scan and the neuronavigation software Localite TMS Navigator 3.4.14 (Localite GmbH, Bonn, Germany). The subject tracker was situated on the forehead and attached using tape and visually monitored throughout the experimental session to ensure stable placement.

#### Transcranial ultrasound stimulation

Transcranial Ultrasound Stimulation (TUS) was delivered using a Sonic Concepts CTX-500 four element transducer (Sonic Concepts Inc, Washington, US) with a 500kHz center frequency, radius of curvature of 6.3cm and an aperture diameter of 6.18cm. Driving system was a Sonic Concepts Transducer Power Output (TPO) (TPO-203-43, firmware version 5.17) through a 4-channel matching network. The verum stimulation target was defined as the grey matter, approximately halfway down the anterior wall of the precentral sulcus at the center of the hand knob (Yousry et al., 1997) in the medioposterior to lateroanterior direction. The target was individualized in the neuro navigation based on each participants anatomy using the t1w image. No noisemasking or earplugs were used during sonication. The active control targets were placed 3cm anterior and posterior to the verum target. The choice of anterior and posterior targets was visually inspected and, if needed, further refined making sure that the chosen target remained within a cortical gyrus.

Stimulation parameters were derived from the tbTUS protocol as described by Zeng et al. (2022) and were set as follows on the TPO: Power/Ch: 3.370W, ISPPA: 9.10W/cm^2,^, ISPTA: 0.91W/cm^2^, Burst length: 20ms, Period: 200ms, Duty cycle: 10%, Duration: 80s. The depth focus of the beam was calculated adding the height of a gelpad (2cm, stardardized) to the scalp-to-target distance as calculated by the neuronavigation software. (2cm) (table 2).

The transducer was attached to a custom 3D printed plastic shell with the purpose of holding a neuro navigation tracker to ensure consistent placement throughout sonication, and a gel pad. Ample amounts of ultrasound gel (Parker Laboratories, INC. New Jersey, U.S.A.) was applied to the transducers active surface and was carefully inspected for bubbles. Bubbles were removed using a small plastic syringe. A 2cm gelpad (Parker Laboratories, INC. New Jersey, U.S.A.) was then placed on the transducers active surface, two different operators visually inspected the gel layer, and air bubbles potentially refracting the focused ultrasound beam, were removed using a plastic syringe. Before coupling the transducer to the scalp, participants hair was parted and a generous amount of ultrasound gel was applied and the hair was gently flattened to fill any voids where air bubbles might reside. Additional gel was then applied and inspected for air bubbles by two different operators, to ensure satisfactory soaking of the hair (Murphy, et al., 2025) The trajectory of the ultrasound beam was assumed be perpendicular to the transducers active surface, and as such the transducer was placed so a straight line could be drawn from the transducer center to the target, while keeping the transducer tangential to scalp, to minimize skull refraction of the beam.

#### Free field acoustic measurements and simulations

Pressure measurements for the transducer were conducted using a 1mm hydrophone (HP1000, Precision Acoustics, UK) with a frequency bandwidth of 0.1MHz to 20MHz assuming an uncertainty of 9% within the frequency bandwidth. The experimental setup included a submersible preamplifier and DC coupler (HP series) connected to a KEYSIGHT 200 MHz digital oscilloscope and robotic arm holding the hydrophone, controlled by a MatLab script developed in-house. The axial and lateral cross-sectional pressure profiles along with corresponding pressure plots were measured for a 9.10 W/cm^2^ ISPPA and 43.5mm focal depth from the transducer exit plane. The free field intensity output of the transducer was measured in a custom water tank (50x30x40cm) filled with deionized water at 22°C. The system was set to produce a continuous sinusoidal wave with a pulse duration of 10ms and pulse repetition interval of 10ms (100% duty cycle). For axial measurements, cross section scans were performed over 60mm x 15mm range with a 1mm step size. For lateral measurements, cross sections scans performed over a 10mm x 10mm range with a 1mm step size (figure 2).

**Figure 1.**
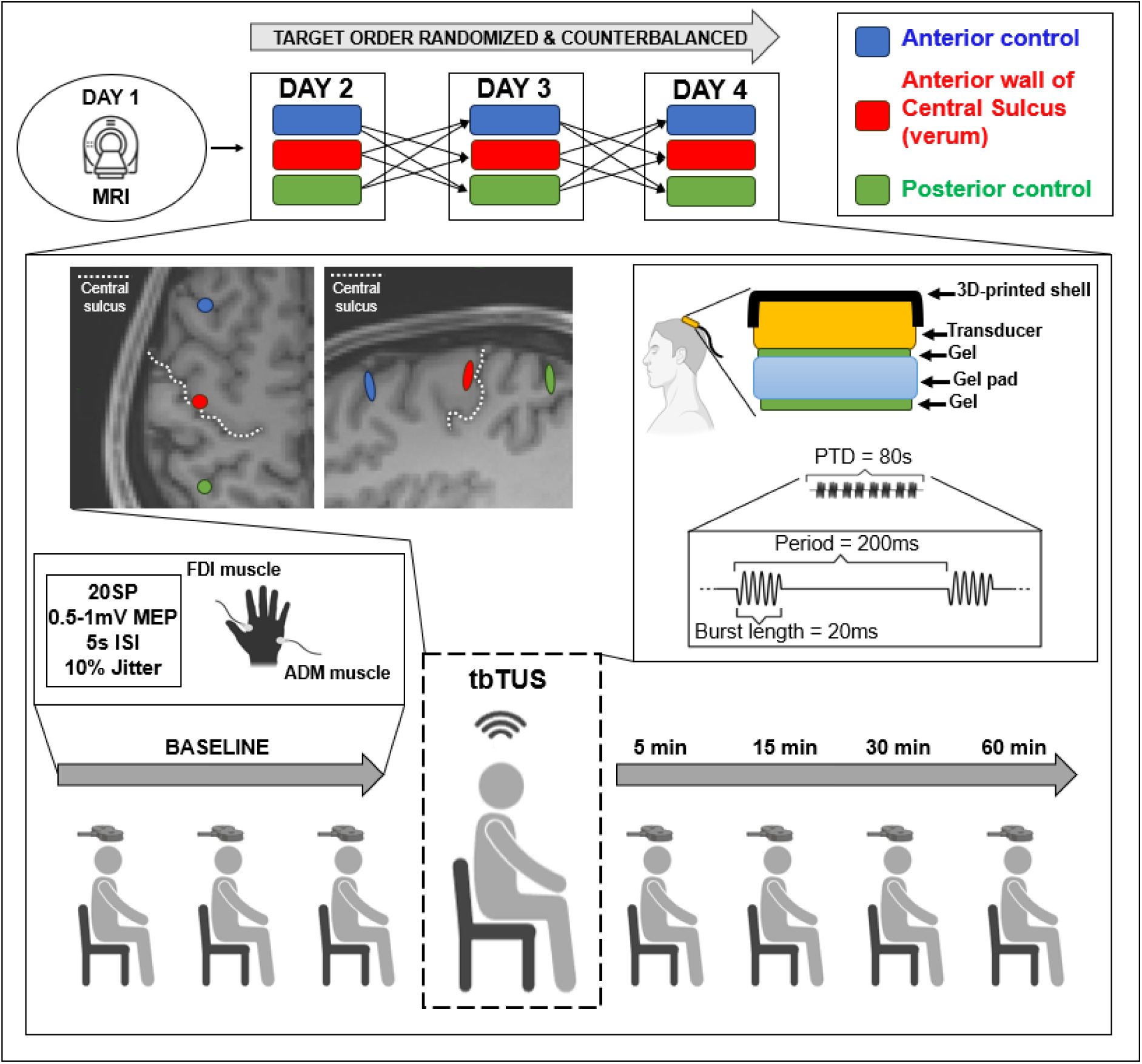
Experimental setup. The experiment was carried out on 4 non-consecutive days. Participants were MR scanned to obtain a T1w image for neuro navigation. For the following three days, tbTUS target order was randomized and counterbalanced to either the anterior wall of Central Sulcus (verum), anterior control or posterior control. Each experimental session was carried out as follows: For the baseline (BASELINE), MEP peak-to-peak amplitudes were measured in 3 blocks using 20 single pulse (SP) TMS delivered to the MEP hotspot with an intensity evoking an MEP with an average amplitude of 0.5 to 1 mV (ISI=5s, jitter ±10%). tbTUS was then applied to the randomized target and follow up MEP measurements were conducted at 5, 15, 30 and 60 minutes after tbTUS (5 min, 15 min, 30 min, 60 min). Coil position was monitored using neuronavigation throughout the entire session. MEPs collected after tbTUS in any condition were not visible to experimenters to avoid potential bias.

**Figure 2.**
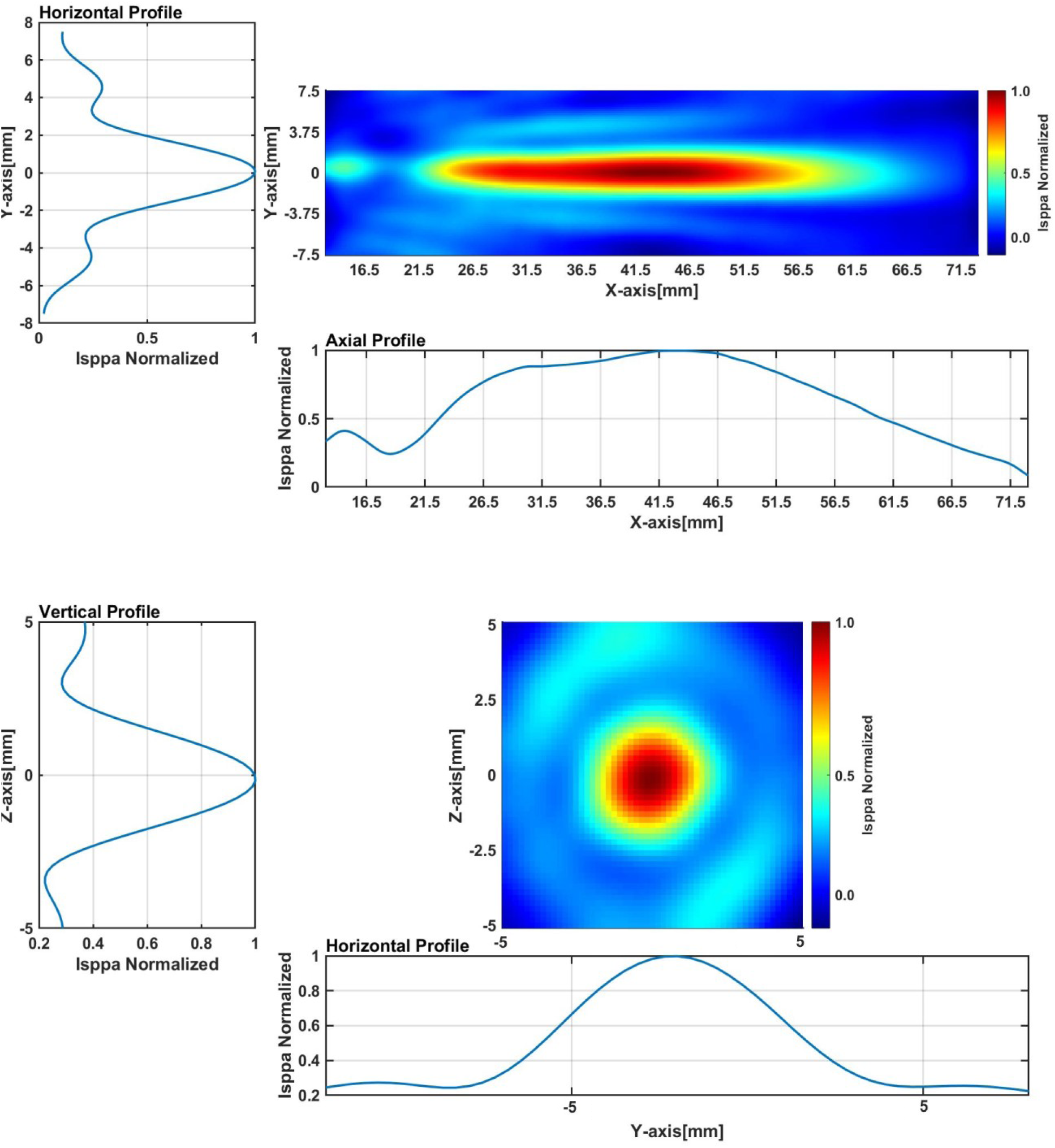
Free field acoustic measurements – The axial (top) and lateral (bottom) cross sections of the measured ultrasound beam and pressure profile for ISPPA = 9.10W/cm^2^, pulse duration = 10ms, pulse repetition interval = 10ms (100% duty cycle) at a focal depth of 43.5mm (Approx. average depth across sessions for verum target).

The oscilloscope’s voltage output was converted into pressure using the following equation: 𝑝 = 𝑉/𝑚(𝑓) where p is the instantaneous acoustic pressure, V is the measured voltage and M(f) the frequency-dependent sensitivity of the hydrophone, and then to intensity using the following equation 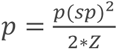 where sp is the instantaneous acoustic focal peak pressure, Z is the acoustic impedance. Intensity was then then normalized on a scale from 0 to 1 and plotted as pressure maps and profiles (Figure 2). From normalized pressure values, the focal width half maximum was calculated from both axial and lateral planes resulting in 32.52mm and 3.32mm respectively. Assuming the focal width of the 3^rd^ dimension is also 3.32mm, the approximated -3db focal volume was calculated, resulting in 398.125mm^3^, which is equal to the -6db normalized pressure amplitude. To obtain the mechanical index (MI), peak pressure, and temperature rise we performed free field acoustic simulations using TUS calculator (Radboud University) using the values reported in Table 1 (appendix). This resulted in a non-derated MI of 0.7674, a peak pressure of 0.54 MPa and a 0.33°C temperature rise. Derating was then applied using TUS calculator (Radboud University) with the following assumptions: 5.8mm scalp thickness, 10mm skull thickness, and an attenuation coefficient of 1.565dB/cm/MHz, resulting in a estimated in-situ pressure amplitude of 0.33MPa and a MI of 0.74. These values (MI, pressure and temperature rise) sit well below the FDA safety guidelines for diagnostic ultrasound and ITRUSST guidelines (Murphy et al., 2025).

**Table 1.**
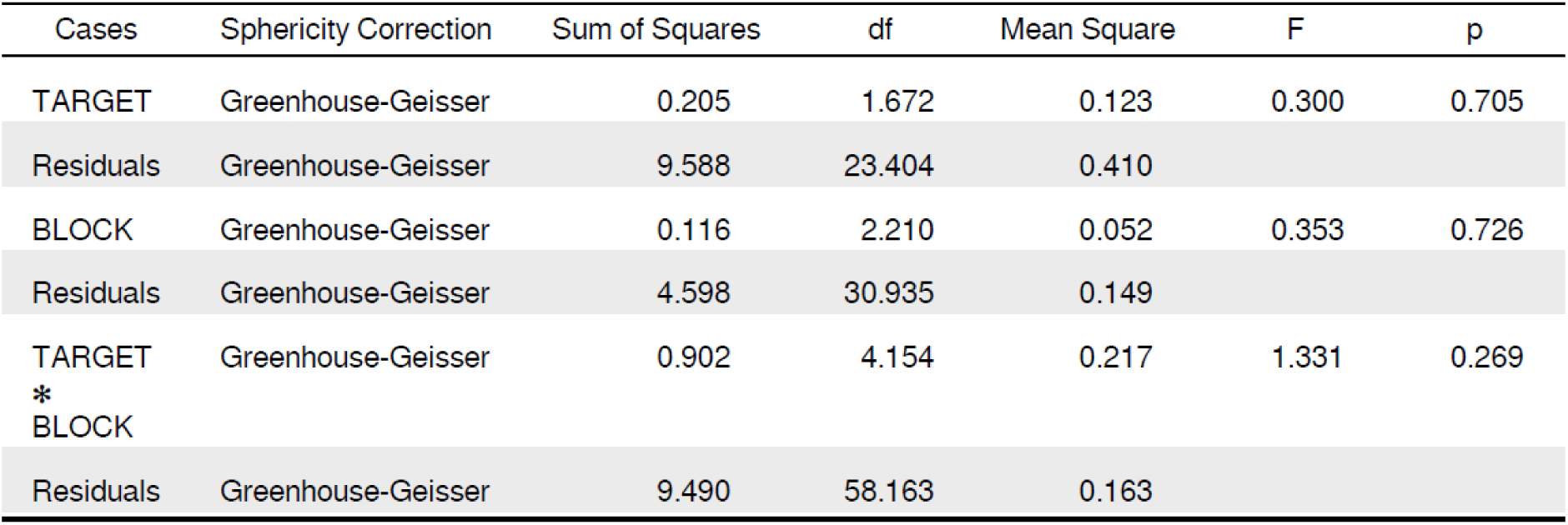
repeated measures ANOVA results.

#### Transcranial magnetic stimulation

Transcranial magnetic stimulation (TMS) was delivered using a MagVenture MagPro stimulator (MagVenture A/S, Farum, Denmark) from a MC-B70 coil with a single biphasic pulse with an AP-PA induced current direction. Participants were seated comfortably in a chair with their hands resting on a pillow in their lap. TMS was delivered to the left M1 hand representation TMS hotspot. The hotspot was defined as the location on the scalp that most consistently elicited MEPs from the FDI hand muscle. After determining the TMS hotspot, stimulation intensity was defined as %MSO that consistently elicited MEPs with an amplitude of 0.5-1mV (SI_0.5-1mV_) from the FDI hand muscle. We ensured that the chosen intensity was sensitive to bidirectional changes by increasing the intensity by a few %MSO and inspecting whether the MEP amplitude increased. This was the case for all the subjects on all days. Stimulations were delivered in blocks of 20 stimulations with an ISI of 5s and 10% jitter. MEPs were acquired at in 3 blocks at baseline interspaced with 5 minutes and in 1 block 5, 15, 30 and 60 minutes after tbTUS.

#### Electromyography

Surface electromyography (EMG) was recorded during TMS from the first dorsal interrosseus (FDI) and abductor digiti minimi (ADM) hand muscles with pairs of surface electrodes (Ambu A/S, Ballerup, Denmark) placed in a belly-tendon montage and a ground electrode placed on the Processus Styloideus Ulna. Before placement, the skin was scrubbed using abrasive gel to ensure optimal signal quality. EMG signal was sampled at 5000hz and amplified with a gain of 500, 2hz lowcut and 2000hz highcut through a Digitimer D360 amplifier (Digitimer Ltd., Welwyn Garden City, United Kingdom) before being converted from analog to digital using a CED 1401 MICRO4 DAC board and visualized using Signal software version 4.11 (Cambridge Electronic Design Ltd., Cambridge, United Kingdom).

#### Data preprocessing

After each session, MEP trials were visually inspected, and trials containing visible background contraction exceeding 25µv, 100ms before the TMS pulse was delivered, was removed from analysis. (mean trials removed: 9 out of 120 (min=0, max=68)).

Before analysis, MEP amplitudes were baseline corrected, to express changes in MEP amplitude following tbTUS as ratios compared to the baseline mean instead of absolute values, as was done in the original work by Zeng et al. (2022).

#### Data analysis

Using JASP (Version 0.19.3) we conducted a repeated measures ANOVA to test for changes in corticospinal excitability following tbTUS at the target sites with TARGET and BLOCK specified as within-subject factors in the repeated-measures ANOVA, ensuring that the crossover structure (each participant completing all TARGET conditions) was modeled appropriately. Greenhouse-Geisser correction was applied in case of non-sphericity. Significance level was set at p<0.05.

Covariation patterns of tbTUS effects across stimulation sites were explored at the within-subject level using baseline-corrected changes in MEP amplitude for each participant from baseline to each of the four post-stimulations, using a pairwise Pearson correlation analysis corrected for multiple comparisons using Benjamini-Hochberg (FDR).

## Results

None of the tbTUS conditions produced group-level changes in MEP amplitude at any time point. The repeated-measures ANOVA revealed no main effect of TARGET (Anterior control, M1, posterior control), BLOCK (5min, 15min, 30min, 60min), and no interaction between the two factors on baseline corrected MEP amplitude (Figure 2) (TARGET: F = 0.300, p = 0.743, DoF = 1.672. BLOCK: F = 0.266, p = 0.899, DoF = 2.210. TARGET x BLOCK: F = 1.158, p = 0.331, DoF = 4.154 (Table 1)).

Independently of target site, changes in corticospinal excitability after tbTUS was highly variable across participants (Figure 2) No consistent changes in MEP amplitude were observed for any of the three target sites relative to baseline (Fig.2).

To assess whether individual tbTUS effects covaried across stimulation sites, we computed baseline-corrected changes in MEP amplitude for each participant from baseline to each of the four post-stimulation time points. For each pair of stimulation sites, these change scores were plotted against one another. We then conducted pairwise Pearson correlation analyses (FDR multiple comparisons corrected) between targets at each time point. Across all comparisons, no systematic relationships or consistent trends were observed (All p>0.05, p = 0.088 – 0.99, r = -0.24 – 0.45 (Figure 3)).

**Figure 3.**
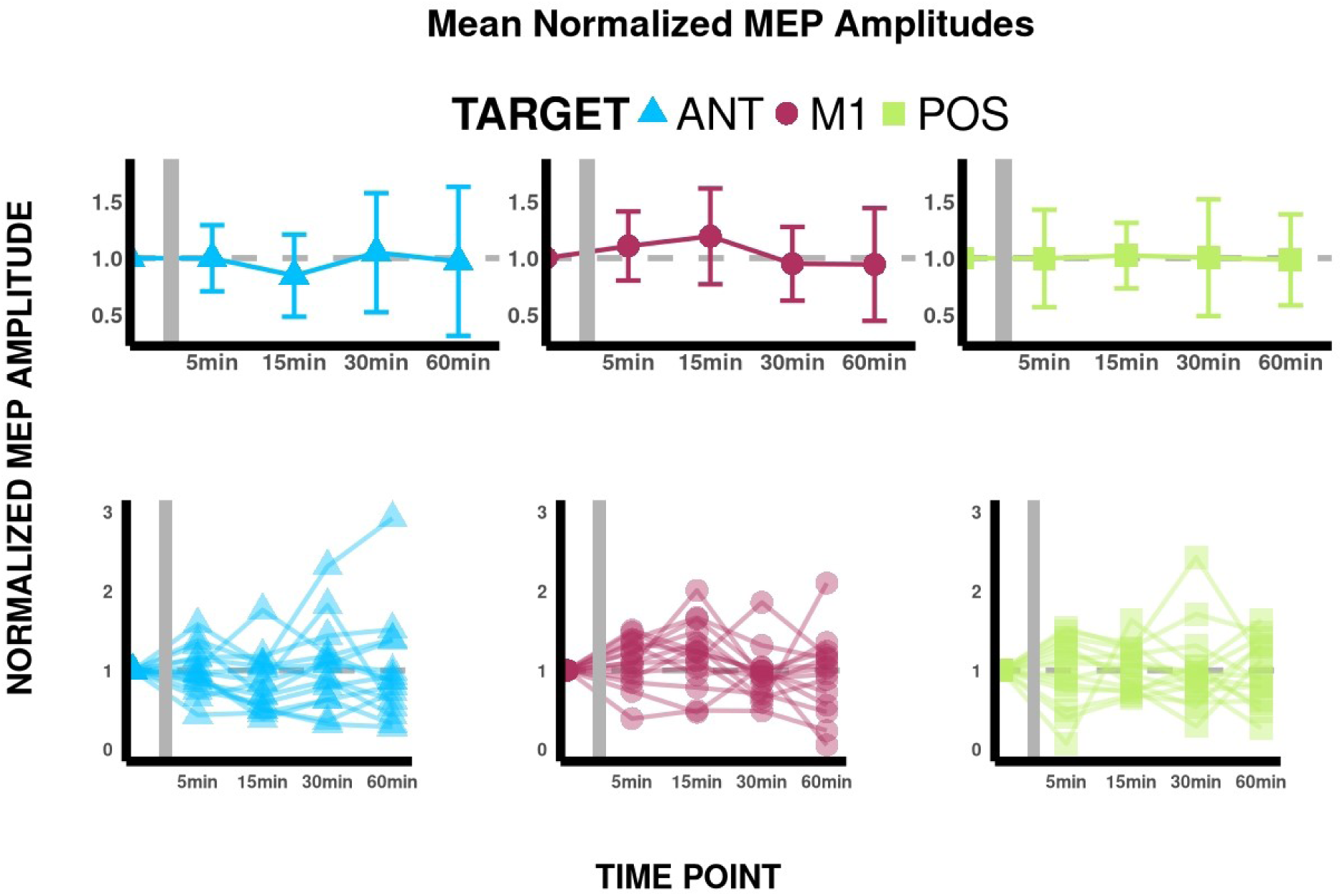
Line plots displaying MEP amplitudes. Top row: Average normalized MEP amplitude for each stimulation site (ANT, M1 and POS), for each time point (Y-axis = baseline, T5 min, 15 min, 30 min and 60 min). Bottom row: Individual average traces, for each participant, for each stimulation site and time point. Error bars represent standard deviation. Grey vertical bar indicates time of tbTUS

**Figure 3.**
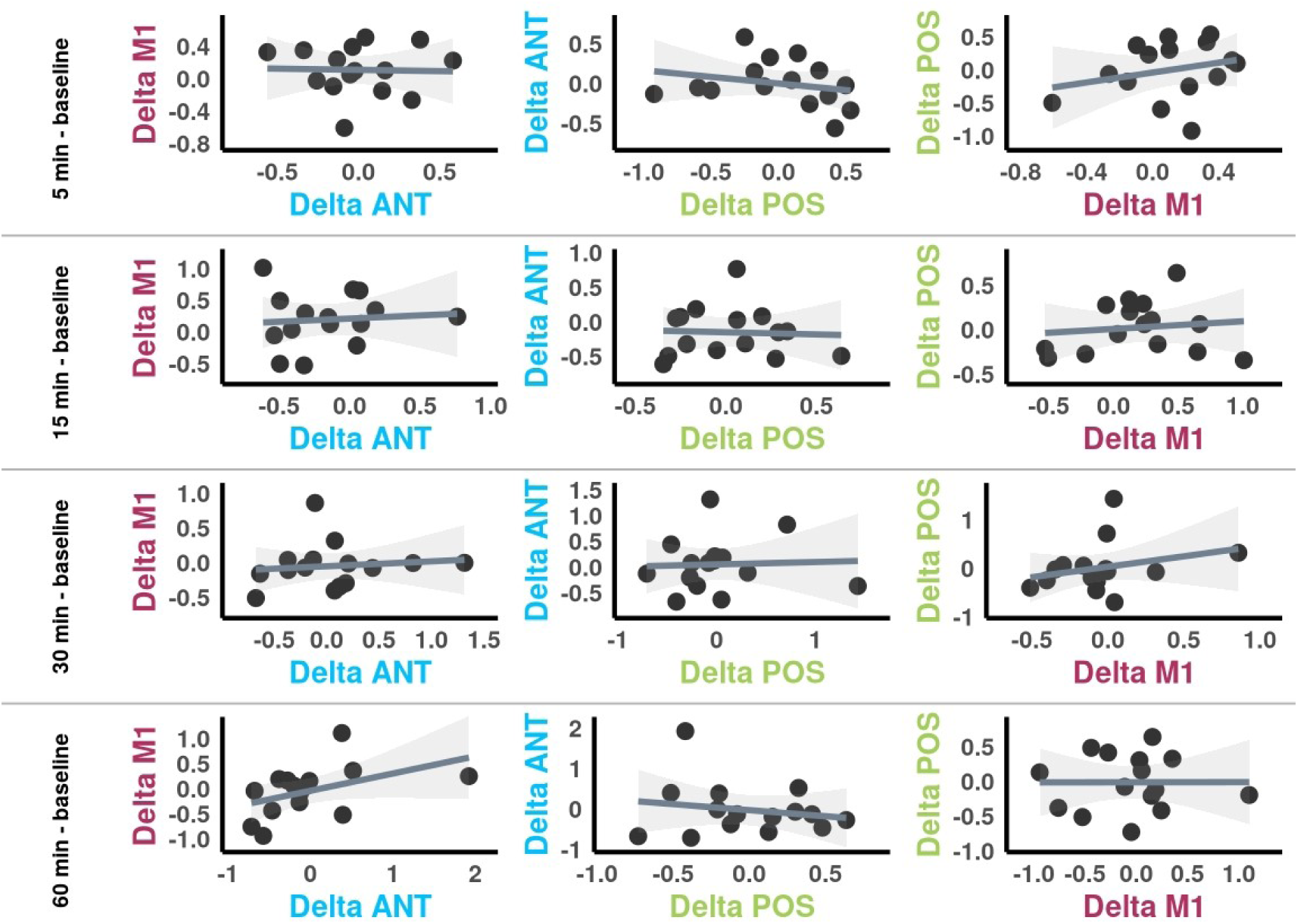
Scatter plots, showing the correlation between delta values for the three conditions, obtained by subtracting normalized MEP amplitudes at 5 min, 15 min, 30 min and 60 min after TUS, with baseline MEP amplitudes.

## Discussion

We found no evidence that tbTUS targeted at the anterior wall of the central sulcus, reliably modulated corticospinal excitability, as indexed by MEP amplitude, relative to control stimulation sites. Individual responses were highly heterogeneous, with MEP amplitudes increasing, decreasing, or remaining unchanged following stimulation, and response magnitudes at one site did not correlate with responses at other sites. This pattern indicates not only an absence of systematic tbTUS effects under the present protocol, but also a lack of stable within-participant responsiveness.

Several factors may account for these null findings. First, our targeting strategy differed from that of Zeng et al. (2022), who used the FDI MEP hotspot as their TUS target, whereas we employed an anatomically defined site on the anterior wall of the central sulcus. These two locations may likely overlap, and the modest discrepancy in sonication target is unlikely to fully explain the absence of group-level changes in the present data.

Second, while we used neuronavigation to position the transducer, we lacked an in-vivo method to verify that the acoustic focus actually reached the intended cortical site. Our approach assumed that a beam perpendicular to the transducer would propagate linearly to the target. However, skull-induced refraction can considerably shift or distort the acoustic field. Fong et al. (2024) demonstrated that transducer placement over the TMS hotspot produced meaningful acoustic overlap with the M1 hand area in only a subset of participants, with >20% overlap in just 33% of their sample. Thus, insufficient or inconsistent target engagement is a plausible contributor to the lack of detectable effects in this study.

Third, the broader tbTUS literature reflects unresolved uncertainty about the reliability, directionality, and boundary conditions of its neuromodulatory effects. Although the original laboratory has reported reproducible facilitation in multiple populations, several independent replications have failed to reproduce these effects. Fong et al. (2024) found no difference between active tbTUS and sham in a double-blind design, even in participants with substantial simulated acoustic-target overlap. Conversely, Bao et al. (2024), who used individualized MRI-based acoustic simulations to ensure precise focus placement, observed sustained inhibition of MEPs relative to sham. These discrepant outcomes, despite rigorous methodologies, suggest that tbTUS effects may depend on factors not yet fully understood—such as cortical state at the time of stimulation, variations in skull morphology, or differential sensitivity of cortical microcircuits.

Beyond these mechanistic considerations, several methodological aspects warrant attention. Our study, like others in this area, may have been underpowered to detect small but reliable tbTUS effects, particularly given the high degree of inter-individual variability. Furthermore, we did not model skull acoustics, quantify acoustic dose at the cortical surface, or incorporate in-vivo validation techniques such as MR-ARFI or ultrasound-sensitive functional MRI (Martin et al., 2025), which could confirm whether the intended tissue was effectively stimulated. It is also worth mentioning that the original work by Zeng et al. (2022) employed a custom made transducer. This is important because the characteristics of a transducer, (eg. number of elements and aperture diameter) directly affects the focality of the ultrasound beam (Murphy et al., 2025). Difference in focality might influence the amount of tissue stimulated, and in turn the after effects. Future work would benefit from integrating these approaches alongside standardized reporting of acoustic parameters to enable meaningful comparisons across studies.

Finally, participant-level moderating variables, such as skull thickness, gyral geometry, baseline corticospinal excitability, and individual neurovascular or neurochemical profiles, may substantially influence responsiveness to tbTUS. Systematically examining these factors will be critical for understanding whether tbTUS has consistent neuromodulatory potential or whether its effects are inherently variable across individuals.

In sum, our findings contribute to a growing body of evidence suggesting that tbTUS effects on human motor cortex are neither robust nor yet mechanistically understood. To advance the field, future studies must combine individualized skull modeling, validated measures of target engagement, harmonized stimulation protocols, and adequately powered designs. Such rigor will be essential to determining whether tbTUS can reliably modulate human cortical excitability and, if so, under what conditions.

## Acknowledgements and conflicts of interest

The authors would like to thank Kim Lara Kürzel for her assistance during data collection and Dr. Xavier Corominas-Teruel for his assistance in free field acoustic measurements, simulations and reporting of intensity parameters.

## Appendix

**Table 2.** TUS Hardware and stimulation parameters.

|  |  |
| --- | --- |
| Hardware |  |
| **Manufacturer** | Sonic Concepts, Neurofus LT, Washington |
| **TPO model #** | TPO-203 S/N: 043 |
| **Transducer model #** | CTX-500_4CH S/N: 038, Ø64 mm x R63.2 mm |
| **Center frequency** | 500kHz |
| **Operating frequency** | 500kHz |
| Stimulation parameters |  |
| **Power/Channel** | 3.370W |
| **ISPPA** | 9.10W/cm<sup>2</sup> |
| **ISPTA** | 0.91W/cm<sup>2</sup> |
| **Burst length** | 20ms |
| **Period** | 200ms |
| **Duty cycle** | 10% |
| **Pulse train duration (PTD)** | 80s |
| **Focus depth in verum target** | Mean = 43.23mm (min: 38.5, max: 49.5) |
| **Focus depth in anterior control target** | Mean = 43.37mm (min: 38, max: 49.5) |
| **Focus depth in posterior control target** | Mean = 42.72mm (min: 38, max: 49.5) |

